# A novel innate lymphoid cell delineates childhood autoimmune arthritis

**DOI:** 10.1101/416784

**Authors:** Martin Del Castillo Velasco-Herrera, Matthew D Young, Felipe A Vieira Braga, Elizabeth C. Rosser, Elena Miranda, Lucy Marshall, Meredyth G LI Wilkinson, Lira Mamanova, Mirjana Efremova, Roser Vento-Tormo, Alex Cagen, Hussein Al-Mossawi, Sarah Teichmann, Adrienne M Flanagan, Lucy Wedderburn, Sam Behjati, Chrysothemis C. Brown

## Abstract

Inflammation in autoimmune disease is mediated by a complex network of interacting cells. Their identity and cross-talk are encoded in messenger RNA (mRNA). Juvenile idiopathic arthritis (JIA), a chronic autoimmune arthritis of childhood, is characterised by synovial inflammation with infiltration of both innate and adaptive immune cells^1^. Activated T cells play a role in disease^2^ but the cell types that drive the recruitment and activation of immune cells within the synovium are not known. Here, we utilised droplet-based and full length single cell mRNA sequencing to obtain a quantitative map of the cellular landscape of JIA. We studied 45,715 cells from the synovial fluid of inflamed knee joints and peripheral blood. We identified a population of synovial innate lymphoid cells (ILCs), shared across patients, that exhibited a unique transcriptional profile in comparison to canonical ILC subtypes. Validation at protein-level across a spectrum of autoimmune arthritides revealed that these ILCs are pathologically expanded in a particular type of JIA. Using statistical tools to assess cellular interactions in synovial fluid, ILCs emerged as a central node of communication, expressing the full repertoire of genes required to orchestrate and maintain the inflammatory milieu. Several ILC-mediated signalling pathways may lend themselves as novel therapeutic targets. Together our findings demonstrate a distinct ILC subtype associated with a tissue-specific childhood autoimmune disease.

---

Children are affected by a range of distinct autoimmune arthritides, collectively termed juvenile idiopathic arthritis (JIA). The cells that mediate these diseases are readily amenable for investigation, when inflamed joints are aspirated to alleviate symptoms. Cells thus obtained have been extensively studied using *in vitro* assays, cell marker analyses and bulk expression profiling^2–4^. Single cell transcriptomics dramatically refines the resolution of biological information that can be gained from synovial fluid aspirates to obtain an unbiased, quantitative readout of the cellular landscape of autoimmune arthritis. This type of analysis has the power to capture previously unidentified, potentially pathogenic cell types^5,6^. Here, we sought to investigate the cellular landscape of JIA at single cell resolution.

Initially we studied the cellular composition of inflamed synovial fluid obtained from two treatment-naïve children suffering from oligoarticular JIA (**Supplementary Table 1**). We performed experiments in a single batch with technical duplicates on the same day (**Figure 1a; Supplementary Table 2**). Immediately after knee joint aspiration, we prepared Ficoll-enriched single cell suspensions of mononuclear cells from synovial fluid and peripheral blood. Count tables of mRNA molecules for each cell were derived using a droplet encapsulation-based method (Chromium 10X^7^; **Supplementary Table 3**).

**Figure 1.**
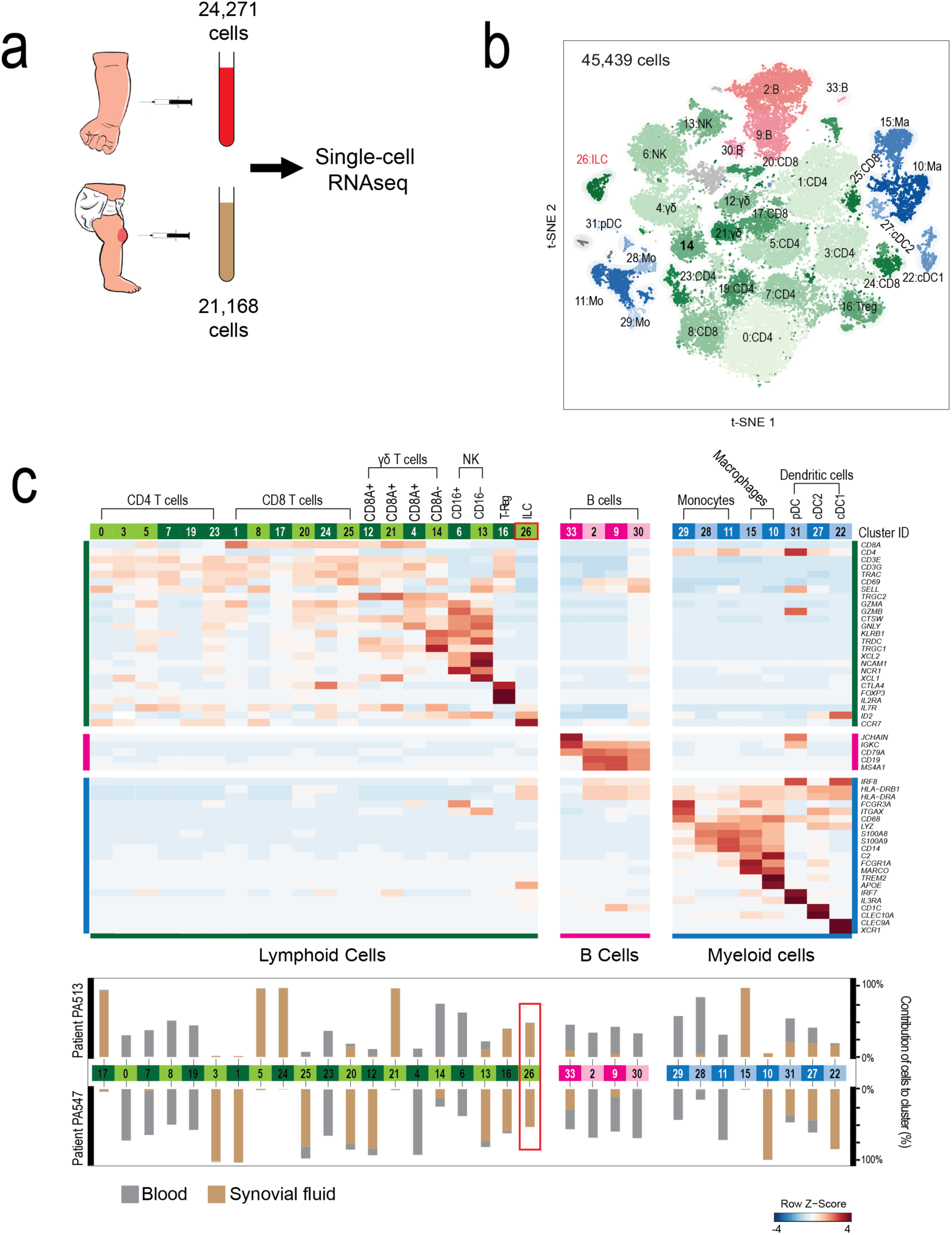
A census of immune cells in childhood autoimmune arthritis. **a.** Overview of experimental design. **b.** Map of cell clusters derived from peripheral blood and synovial fluid cells using t-SNE. Number next to each cluster refers to annotation as per 1C. Clusters that could not be un-ambiguously identified (18 and 32) were removed from further analyses. **c.** Signatures of immune cell types. The relative expression of marker genes (rows) across cell clusters (columns) is shown. Lower panel shows composition of each cluster by tissue and patient. Red square highlights ILC cluster.

Using a community detection algorithm, we derived a reference map of 45,349 cells (**Figure 1b**). We assigned a cellular identity to each cluster by cross-referencing cluster-defining transcripts with canonical markers (**Figure 1c, Supplementary Table 4**). All clusters exhibited the pan-immune cell marker *PTPRC* (CD45, **Extended Data Fig. 1a**) and represented the spectrum of known myeloid and lymphoid cells: CD8, CD4, Treg, γδ T cells, NK cells, B cells, monocytes, macrophages, conventional and plasmacytoid dendritic cells (DC). Synovial fluid immune cells were enriched relative to peripheral blood (p<10^−4^; Fisher’s exact test) for CD8^+^ T cells, regulatory T cells (Treg) and macrophages (**Extended Data Fig. 1b**). These synovial cells were transcriptionally distinct from analogous cell types in peripheral blood.

A striking finding was a cluster of patient-overarching synovial cells (cluster 26, **Figure 1 b, c**) which exhibited a profile that defines innate lymphoid cells (ILC)^8^: negative for leukocyte lineage markers whilst expressing *IL7R* and *ID2* (**Figure 1c**). As a unifying cell population exclusive to synovial fluid, these ILCs (hereinafter referred to as JIA ILC) represented a plausible cell population that may be pathogenic. Canonical ILCs have previously been detected in synovial fluid of patients with rheumatoid arthritis (RA)^9–11^. Intriguingly, the ILCs we identified here lacked expression of lineage defining genes of canonical ILC subtypes^12^ (**Extended Data Fig. 1c**).

To validate JIA ILCs, we devised a panel of markers for flow-cytometry that we used to confirm the presence of JIA ILC in synovial fluid (**Figure 2a**) and show their absence in blood (**Extended Data Fig. 2a**). Cytological analysis of ILCs isolated by FACS confirmed lymphoid morphology (**Figure 2b**). Next, we investigated whether synovial ILCs were a universal feature of the cellular landscape of inflamed joints, or whether they delineated a specific disease. We studied synovial fluid from 32 children, at diagnosis or at relapse following treatment, with different JIA phenotypes (**Supplementary Table 1**): oligoarticular JIA, polyarticular rheumatoid factor negative JIA, systemic JIA and psoriatic JIA. Flow-cytometric assessment of the joint fluid, revealed significant enrichment (p<0.01 and p<0.05, Mann-Whitney *U* test) in two disease phenotypes: oligoarticular JIA and polyarticular rheumatoid factor negative JIA, considered to represent the same disease^13^ (**Figure 2c**). This would suggest that JIA ILCs characterise a specific autoimmune disease of childhood.

**Figure 2.**
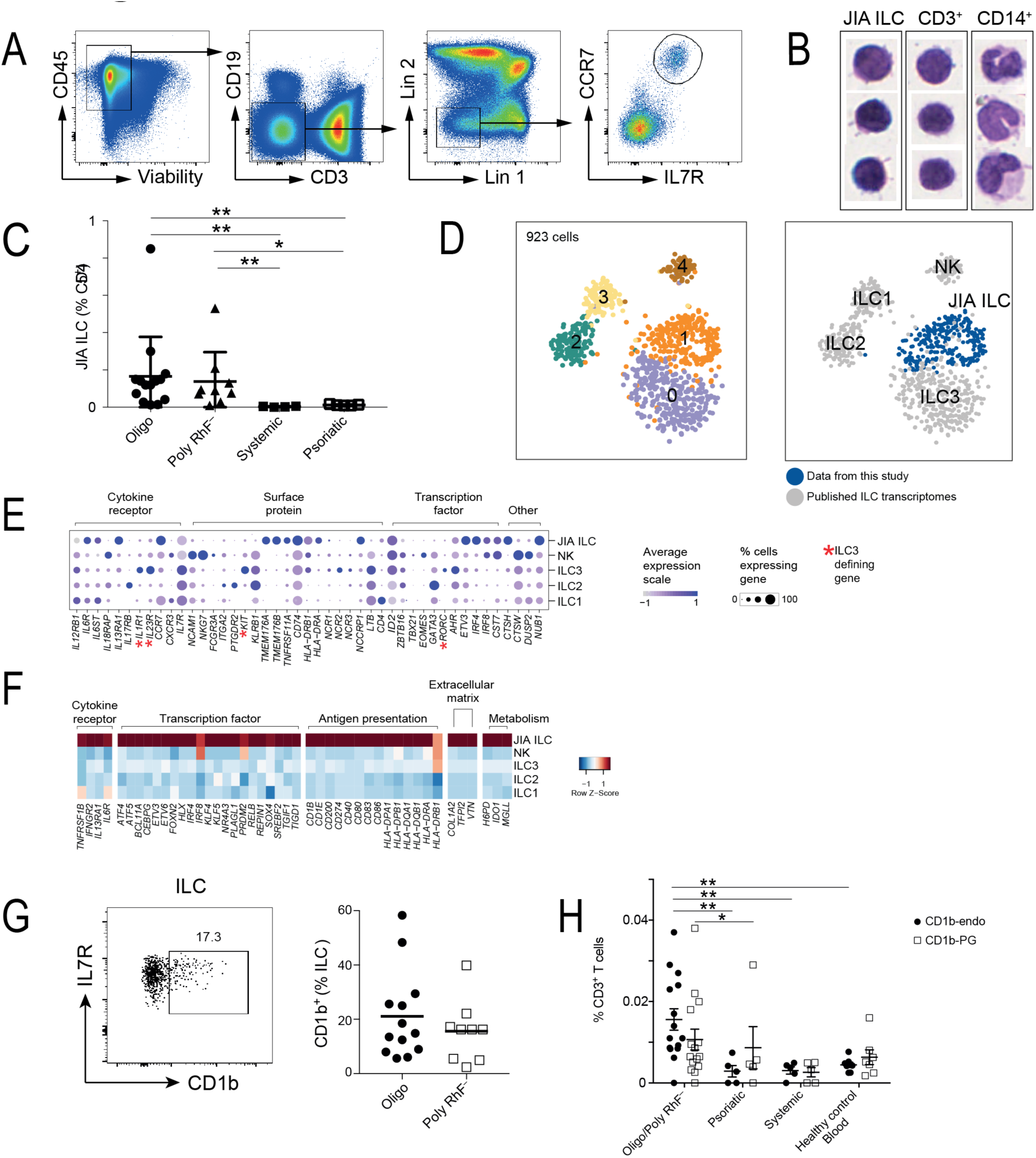
Characterisation of JIA ILCs. **a.** Gating strategy for identification of synovial ILCs by flow cytometry with JIA ILCs defined as Lin (CD34, CD14, CD16, CD56, FceR1a, CD123, CD11c, XCR1, CLEC9A, CD3, CD19)-CD45+IL7R+CCR7+. **b.** Morphology of JIA ILCs, CD3+ and CD14+ cells sorted from synovial fluid. Cells were fixed and stained with Giemsa. **c.** Frequency of JIA ILCs (Lin-CD45+IL7R+CCR7+) as a percentage of CD45+ cells in synovial fluid from patients with distinct autoimmune diseases affecting the joint: oligoarticular JIA (Oligo), polyarticular rheumatoid factor negative JIA (Poly), systemic JIA (Systemic) and psoriatic JIA (Psoriatic), (mean □ SEM; * P<0.05, ** P<0.01, Mann-Whitney U test) **d.** t-SNE of 276 JIA ILCs and 647 human ILCs from tonsils, coloured by subcluster (left panel) or synovial fluid ILCs (right panel). **e.** Expression of ILC-related genes across known ILC subsets and JIA ILCs **f.** Heatmaps of average gene expression, standardized by z-scores, across all cells within each cluster (JIA ILC, ILC1, ILC2, ILC3, NK cells) for cytokine receptors, transcription factors, antigen-presentation molecules and extracellular matrix remodeling proteins. **g.** Representative flow cytometry dot plot showing cell-surface expression of CD1b on JIA ILCs. Number indicates percent CD1b-pos itive cells (left panel). Right panel shows summary of frequency of CD1b+ JIA ILCs from patients with oligoarticular (Oligo) or polyar ticular rheumatoid factor negative JIA (Poly). **h.** Frequency of CD1b-endo or CD1b-PG tetramer positive cells in the CD3+ population in synovial fluid of patients with different childhood autoimmune arthritides: oligoarticular JIA (Oligo), polyarticular rheumatoid factor negative JIA (Poly RhF-), Systemic JIA (Systemic) and Psoriatic JIA (Psoriatic) or blood from healthy paediatric donors (healthy control); (mean □ SEM; * P<0.05, ** P<0.01, Mann-Whitney U test)

To gain further insight into the identity and function of JIA ILCs we profiled full-length transcriptomes of 276 JIA ILCs using smart-seq2 scRNA-seq (**Supplementary Table 5**). We obtained cells from frozen synovial fluid samples of one of the initial children and one additional child (PA400) presenting with the same disease phenotype (**Supplementary Table 1**). The transcriptional profile of JIA ILCs thus obtained aligned with the JIA ILCs identified in the initial orthogonal experiment (**Extended Data Fig 3a**). Comparison of JIA ILC transcriptomes to canonical ILCs isolated from human tonsils^14^ revealed an independent cell cluster, largely (87%) composed of JIA ILCs (cluster 1, **Figure 2d)**. JIA ILCs lacked transcripts that characterise canonical ILC subsets including genes that define the ILC3 lineage (*RORC, IL23R, IL1R1* and *KIT*; **Figure 2e**). However, JIA ILCs shared expression of some genes, beyond *IL7R* and *ID2*, with other ILCs, such as genes implicated in the identity and function of ILC3s (e.g *AHR, TNFRSF11A, TMEM176A TMEM176B*^12,15,16^; **Figure 2e**).

**Figure 3.**
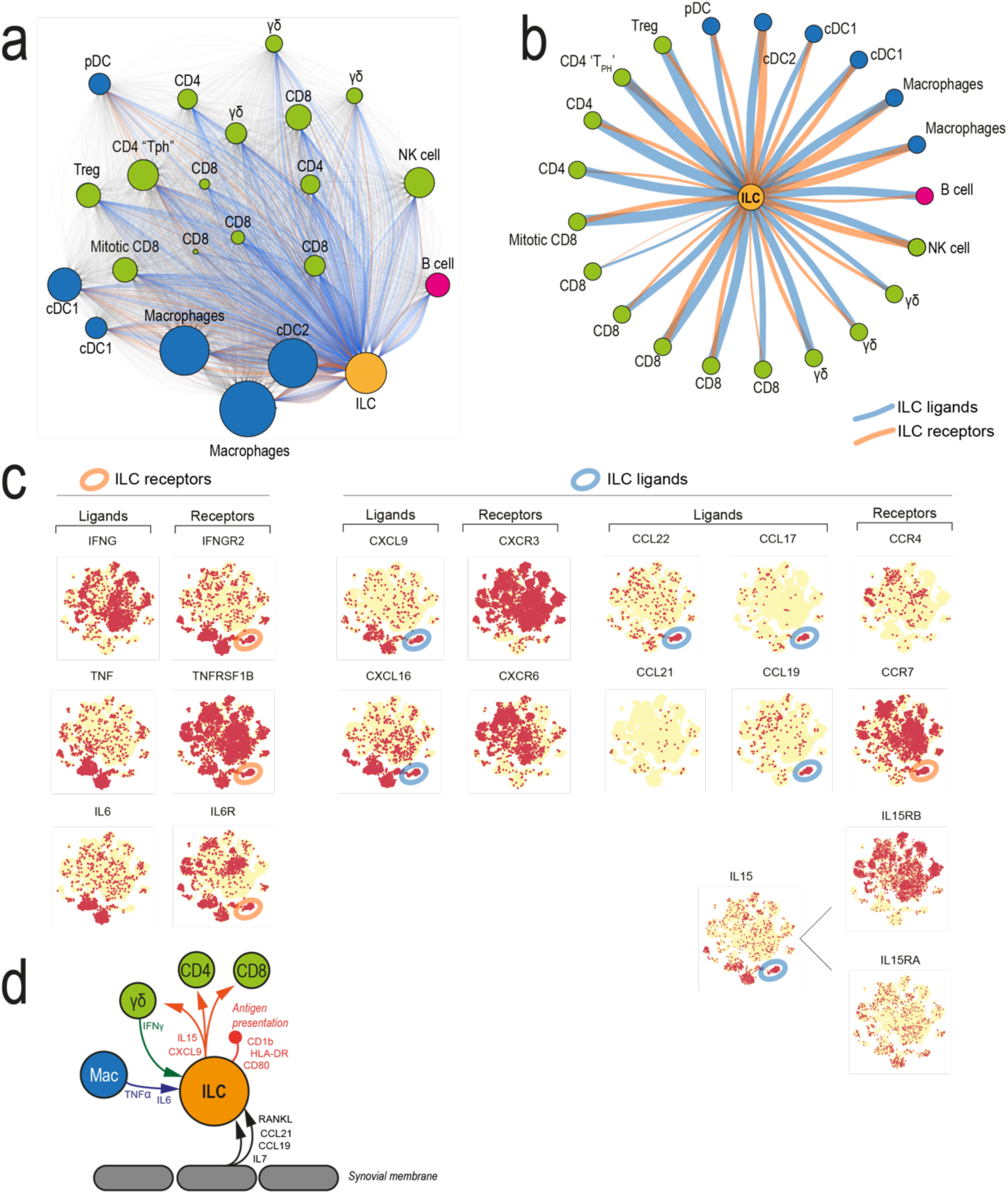
Analysis of cellular interactions within joint exudates. **a.** Overview of significant receptor/ligand interactions in the synovial fluid of both patients. Each circle is a cluster of cells, and each line a receptor/ligand pair significantly expressed by the connected clusters in one patient (see methods). ILC receptors are shown in blue, ILC ligands are showing in red and other receptor/ligand pairs in grey with low opacity. Cluster sizes are proportional to the total significant receptor/ligand count, excluding self-interacting pairs. **b.** Summary of the interactions between JIA ILCs and all other clusters. ILC receptors/ligands are shown in red/blue respectively with the line thickness proportional to the total number of connections to each cell type. **c**. Gene expression for ILC cytokine receptors, chemokines, cytokines and corresponding ligands/receptors, overlaid on the t-SNE visualization (Extended Data Figure 5a). Yellow indicates absence of expression within a cell, red indicates presence of transcript. The ILC cluster has been circled in blue where the ligand is expressed highly and in orange when the receptor is expressed highly. **d.** Schematic illustration of JIA ILC signaling.

JIA ILCs exhibited a unique ILC phenotype in terms of transcription factors and cytokine receptors (**Figure 2f, Extended Data Fig. 3b**). The transcriptional profile of JIA ILCs was further distinguished by expression of genes associated with extracellular matrix remodelling (*COL1A2, VTN, TFPI2*; **Figure 2f**, **Extended Data Fig. 3b**) indicating a possible contribution of synovial ILCs to tissue homeostasis. JIA ILCs did not express canonical markers of osteoclasts or synovial fibroblasts (**Extended Data Fig. 3c**). Furthermore, JIA ILCs were notable for expression of transcripts related to antigen presentation (eg, HLA class II genes, *CD74, CD1B, CD1E*) and leukocyte activation (*CD80, CD86, CD40, CD83;* (**Figure 2e, f**, **Extended Data Fig. 3b**).

Whilst subsets of ILCs are known to express HLA/MHC genes^17,18^, the specific profile of JIA ILCs with expression of the antigen presenting molecule, *CD1B*, and the lipid antigen loading molecule *CD1E*, along with co-stimulatory molecules, suggest that JIA ILCs possess the complete repertoire for T cell priming through both peptide and lipid-antigen presentation. *CD1B* expression is of particular interest because its expression on ILCs was previously unknown and expression of another CD1 family member, CD1a, has been detected in skin ILCs in response to inflammatory signals^19^. We confirmed the presence of CD1b on synovial fluid ILCs (**Figure 2g**) and assessed the frequency of autoreactive CD1b-restricted T cells across a range of paediatric synovial pathologies using CD1b tetramers loaded with endogenous lipid (CD1b-endo) or phosphatidylglycerol (CD1b-PG) (**Figure 2h, Extended data Fig. 4a, b**). The proportion of CD1b-reactive T cells was significantly higher in the synovial fluid of children with JIA (P<0.01 in all comparisons with CD1b-endo; Mann-Whitney *U* test, **Figure 2h**) compared to other arthritides or blood from healthy children. Given that these two tetramers identify CD1b-reactive T cells specific for a fraction of potential CD1b binding antigens^20,21^, it is possible that a greater proportion of synovial T cells are CD1b-restricted. A highly upregulated gene in synovial ILCs was the monoglyceride lipase-encoding *MGLL* (**Figure 2f, Extended Data Fig. 3b**). Increased *MGLL* expression has previously been reported to generate an altered profile of free fatty-acids, mono-acyl glycerol and secondary lipid metabolites^22^. We speculate that disordered lipid metabolism coupled with CD1b expression leads to altered presentation of lipid antigens to self-reactive T cells within the synovium. Interestingly, previous studies have shown that ILCs exhibiting CD1 molecules are able to prime T-cell responses *in vitro*^19^.

Assessment of JIA ILCs by additional *in vitro* assays was not feasible given the small numbers of ILCs found in inflamed joint. Therefore, we sought an alternative approach for studying ILC function within the context of the synovial inflammatory milieu, *quasi in situ.* We analysed the cellular communication network across the clusters present in each joint (**Extended Data Figure 5a, b**). We determined statistically significant interactions between clusters, defined as expression of receptor-ligand pairs, using the CellPhoneDB algorithm (Vento-Tormo R *et al.*; in press in *Nature*)^23^. Judging by the number of cellular interactions, ILCs emerged as a central communication node within the autoimmune infiltrate of each child (**Figure 3a**). ILCs possessed a broad array of cytokine receptors (*TNFRSF1B, IL6R, IFNGR2, IL27RA, IL3RA1*; **Figure 2f, 3c**) allowing detection of macrophage and DC derived cytokines, as well as the full complement of chemokines required for recruitment of all effector subsets (*CXCL9, CCL22, CCL17*; **Figure 3c**). Notable amongst effector cells were a cluster of actively dividing MKI67^+^ CD8 T cells (cluster 12, **Extended Data Fig 5c)**, as well as a cluster of CD4 T-cells expressing *CXCR6, CCR5, PDCD1, MAF* and *ICOS* (cluster 10, **Extended Data Fig 5d**); a gene profile previously reported on RA synovial ‘T_PH_’ cells^24,25^. Central to ILC-T cell interactions may be ILC-derived IL-15 (**Figure 3c)**. IL-15 has been previously identified in inflamed synovial fluid^26^ and drives expansion and maintenance of memory T-cells and NK cells^27^. Although stromal-ILC interactions were not captured in our experiment, we note high levels of *CCR7* and *TNFRSF11A* (RANK) in JIA ILCs (**Figure 2e, 3c**), enabling communication with stromally derived CCL19/21 and RANKL^28,29^. Together these analyses define the cross-talk within the synovial inflammatory exudate, including specific interactions that may be amenable to therapeutic intervention with existing drugs, such as inhibition of IL15 or augmentation of RANK signalling.

Finally, we integrated gene expression profiles of synovial cell types with candidate risk genes from genome-wide association studies (GWAS) in JIA patients^30,31^. This revealed prominent expression in ILCs of *TRAF1* and *ZFP36L1* (**Extended data Fig 6a**), further supporting a functional role for ILC-mediated cytokine signalling and production in disease pathogenesis.

In this investigation we identified a novel ILC subtype that is pathologically expanded in a specific childhood autoimmune arthritis. Although it is currently not feasible to prove directly that synovial joint ILCs drive JIA, collectively our findings are consistent with a model of synovial ILCs contributing to, if not primarily orchestrating, the aberrant immune response of JIA joint inflammation. ILCs are largely tissue-resident cells that seed tissues during foetal and early life, acquiring organ-specific functions^32^. The expression of tissue-context specific genes by synovial ILCs raise the possibility that ILCs are in fact resident in synovial tissue and become aberrantly activated in JIA. Thus, the cellular roots of non-systemic JIA, a disease with defined onset in childhood, may lie in human development. Given that JIA shares many disease-associated risk alleles with other childhood autoimmune diseases^33^, including type 1 diabetes and IBD, we speculate that the cellular and molecular pathways revealed by our analyses may be pertinent to other tissue-specific autoimmune diseases.

## Methods Summary

### Study subjects

Blood and synovial fluid samples were collected from JIA patients undergoing intra-articular aspiration at Great Ormond Street Hospital, London. Healthy child and adolescent peripheral blood samples were obtained from the Arthritis Research UK Centre for Adolescent Rheumatology at UCL, UCLH and GOSH Biobank. The study was performed in accordance with protocols approved by the Bloomsbury Research Ethics Committee (London, UK) and North Harrow Ethics Committee. Full informed consent was obtained from patients and/or parents.

### Cell isolation

Synovial fluid was treated with hyaluronidase (Sigma-Aldrich) at 10 units/ml for 30 min at 37?°C Synovial fluid and peripheral blood mononuclear cells (SFMCs and PBMCs) were then isolated by density-gradient centrifugation using Lymphoprep (Alere). Cells were either analysed immediately (droplet based single cell RNA sequencing) or cryopreserved in freezing media (90 % heat inactivated fetal calf serum (FCS), 10% DMSO) and stored in liquid nitrogen.

### Flow cytometry staining and cell sorting

Single cell suspensions of thawed SFMCs and PBMCs were stained with viability dye (Ghost Dye Violet 510, Tonbo) in PBS followed by a panel of fluorescently conjugated antibodies in staining buffer (PBS, 2mM EDTA, 2% FCS). The following antibodies were used for defining lineage marker cells: PE-CD14 (HCD14, Biolegend); PE-CD16 (B73.1, Biolegend); PE-CD11c (3.9, BD); PE-XCR1 (S15046E, Biolegend); PE-CLEC9A (8F9, Biolegend); Percp-Fc*ε*R1*α* (AER37, Biolegend); Percp-Cy5.5-CD34 (581, Biolegend), Percp-Cy5.5-CD123 (6H6, Biolegend), Percp-Cy5.5-CD56 (HCD56, Biolegend); AF700-CD19 (HIB19, Biolegend), BV711-CD3 (UCHT1, Biolegend). Additional antibodies used for ILC identification included APC-e780-CD45 (H130, Ebioscience); eFluor450-CD127 (eBioRDR5, Ebioscience); PE-Cy7-CCR7 (G043H7, Biolegend); FITC-CD1b (SN13, Biolegend). Flow cytometry and FACS sorting were performed on a BD LSR II or FACS Aria. Data were analyzed using FlowJo (Tree Star, Inc.) software.

### CD1b Tetramers

CD1b monomers were produced in HEK293 human embryonic kidney cells (NIH Tetramer Facility). CD1b monomers were used without additional loading or loaded with phosphatidylglycerol (PG, Avanti Lipids). For loading of CD1b monomers, 32 μg of PG was sonicated for 2 hours at 37?°C in 90 μl of 0.5% CHAPS and 50?mM sodium citrate buffer, pH 7.4. CD1b monomers (20 μg) were added to the tubes and incubated overnight at 37?°C. The following day, PBS was added to reach a final concentration of 0.2?mg/ml loaded CD1b monomers. Streptavidin-PE (Invitrogen) was added to CD1b monomers (CD1b-endo) or PG-loaded monomers (CD1b-PG) for tetramerization. Jurkat 76 (J76) T cells, lacking endogenous TCR α and β chains ^34^, transduced with the PG90 TCR specific for CD1b-PG (J76.PG90)^20^ were used to confirm specificity of CD1b tetramer staining. For identification of CD1b-reactive T cells in patients and healthy controls, thawed SFMCs and PBMCs were first stained with viability dye (Ghost Dye Violet 510, Tonbo) in PBS. Cells were then washed and stained with tetramer in staining buffer (1:50 dilution) at RT for 20 min followed by staining with a panel of fluorescently conjugated antibodies (Percp-Cy5.5-CD3 (OKT3, Biolegend); FITC-TCRγδD(B1, Biolegend); AF700-CD19 (HUB19, Biolegend); BV605-CD14 (M5E2, Biolegend); APC-CD8 (SK1, BD); BV711-CD4 (SK3, BD).

### Cytospin and immunostaining

Cytospin preparations were carried out by centrifugation of FACS-purified CD3^+^, CD14^+^ cells and ILCs at 600rpm for 5 minutes in a Cytospin™ 4 Cytocentrifuge (ThermoFisher Scientific). Slides were fixed for 1 minute in 10% formalin buffer (Sigma) and stained with Giemsa.

### Droplet based single cell RNA sequencing

Cell suspensions from peripheral blood and synovial fluid for two patients with oligoarticular juvenile idiopathic arthritis were prepared. Subsequently, cells were counted using a Neubauer haemocytometer. To obtain technical replicates, each cell suspension was split and loaded into two wells of the Chromium controller (10X-Genomics) chip following the standard protocol for the Chromium single cell 3’ Kit v2 (10X Genomics). A cell concentration was used to obtain an expected number of captured cells between 5000-10000K cells. All subsequent steps were done based on the standard manufacturers protocol. Each 10X chip position was sequenced across a single Illumina HiSeq 4000 lane.

### Plate based single cell RNA-sequencing

Single cells from two patients were individually flow sorted into 96-well full-skirted plates (Eppendorf) containing 10µL of a 2% Dithiothreitol (DTT, 2M Sigma-Aldrich), RTL lysis buffer (Qiagen) solution. To mitigate batch effects, the cells of the different patients were sorted into plates following a balanced blocked design (alternating rows within the same plate). Cell lysates were sealed, mixed and spun down before storing at −80 °C. Paired-end multiplexed sequencing libraries were prepared following the Smart-seq2 (SS2) protocol^35^ using the Nextera XT DNA library prep kit (Illumina). A pool of barcoded libraries from four different plates were sequenced across two lanes on the Illumina HiSeq 2500.

### Droplet based single cell RNA-seq data mapping and pre-processing

The raw single-cell sequencing data was mapped and quantified using the 10X Genomics Inc. software package CellRanger (v2.0.2) against the GRCh38 reference provided by 10X with that release. Using the table of unique molecular identifiers produced by Cell Ranger, we identified droplets that contained cells from empty droplets by two different methods: the EmptyDrops package (github.com/TimothyTickle/hca-jamboree-cell-identification) which finds droplets that vary significantly to the profiles observed from empty droplets (<100 UMIs); and the call of functional droplets done by Cell Ranger. We retained for further analysis all droplets that either were identified by EmptyDrops as significantly different from the background with and FDR <0.01 or called by Cell Ranger with the default parameters as droplets containing a cell.

### Quality control of droplet based single cell data

After we identified cell containing droplets, we proceeded to filter out lower quality cells by removing all cells that showed a percentage of expression from mitochondrial genes higher than 5%; and cells that expressed less than 200 genes. Finally, in an effort to remove droplets that captured two cells (doublets) we removed cells that showed a number of expressed genes higher than 2940 genes, 2.5 standard deviations higher than the average number of expressed genes across the entire droplet-based dataset.

### Background contamination correction in droplet based single cell data

A previous report has showed that background contamination in droplet based single cell data can have serious effects in the biological interpretation of the data (www.biorxiv.org/content/early/2018/04/20/303727). In an effort to reduce the effects of cell free RNA, we used the package SoupX (v0.2.2, www.biorxiv.org/content/early/2018/04/20/303727) to quantify the background contamination and estimate corrected expression profiles. To estimate the background contamination fraction present in each individual 10X channel, we used either a list of haemoglobin genes or immunoglobulin genes (**Extended data table 8)** along with the raw UMI count matrices and the barcodes of the cells called by EmptyDrops and cell ranger. Finally, using the calculated background contamination fraction we generated a table of corrected UMI counts which was used for all subsequent analyses.

### Peripheral blood and synovial fluid droplet based single cell data combined analysis

Having obtained the background corrected table of counts, downstream analyses were done using R (3.4.1) and the Seurat package (v 2.3.0, satijalab.org/seurat). All the blood and synovial fluid data was normalised and log transformed using the “NormalizeData” function with default parameters. Subsequently, we scaled the data using the ScaleData function providing the total number of UMIs and the mitochondrial fraction as factors to regress out. Afterwards, we identified variable genes using the function “FindVariableGenes with the following parameters (x.low.cutoff = 0.0125, x.high.cutoff = 3, y.cutoff = 0.5). Then, using the resulting list of variable genes we performed a principal component analysis considering the first 30 PCs. Subsequently, we identified clusters using a k-nearest neighbour algorithm based on the first 30 PCs and a resolution of 1.2. Finally, we built a two-dimensional visualisation of the data using the t-distributed stochastic neighbor embedding (*t*-SNE) algorithm using the first 30 PCs. Clusters were annotated manually using canonical markers. Clusters with ambiguous identity were excluded from downstream analyses.

In addition to the aforementioned analysis, we performed an independent analysis looking solely at the synovial fluid data. This was carried out following the same workflow as above with the following changes: the clustering was done using the first 28 PCs and a resolution parameter of one. Similarly, the *t*-SNE visualisation was generated using the first 28 PCs. Cluster identity was assigned manually based on known markers.

### Synovial fluid ILC plate based single cell RNA-sequencing expression quantification

To assess the expression from the SS2 data, raw reads were pseudo-mapped and counted using the using kallisto v0.43.1^36^ based on the annotation made by ENSEMBL(v90) of the human reference genome (GRCh38). To obtain per gene counts, all the transcript counts were summarised using scater v1.6.3^37^. Cells with more than 10% of their mRNA originating from mitochondrial genes, a total number of reads <500,000 or a total number of counts >3 median absolute deviations (MAD) were filtered out prior to downstream analysis.

### Comparison of plate-based JIA ILCs single cell data with published single cell ILC data

To compare the expression profile of the JIA ILCs with canonical ILC subsets we utilised publically available single cell expression profiles for human tonsillar NK cells, ILC1, ILC2 and ILC3s, generated using the SMART-seq2 protocol (European Nucleotide Archive (ENA) project ID PRJNA289086)^14^. FASTQ files for a total of 647 cells were downloaded. To make the datasets comparable, the raw sequencing was reanalysed using the same parameters applied to the analysis of SS2 data for JIA ILC. Counts were generated using kallisto v0.43.1^36^ based on ENSEMBL’s annotation(v90) of the human reference genome (GRCh38). To obtain per gene counts, all the transcript counts were summarised using scater v1.6.3^37^. All cells that had 10% of their expression originating from mitochondrial genes were filtered out.

The published ILC gene counts in conjunction with the JIA ILC data were loaded into R computing environment (3.4.1) and processed using the Seurat package (v 2.3.0, **http://satijalab.org/seurat/**). Downstream analysis such as data normalisation and identification of variable genes were run with default parameters. Next, we scaled all the data using the ScaleData function providing the total number of counts, mitochondrial fraction, project and patient of origin as variables to regress out. Subsequently, with the variable genes we performed a principal component analysis using the first 20 PCs. Afterwards, we used the same PCs to build a two-dimensional *t*-SNE representation of the data using a perplexity of 30. Clusters of cells were then identified using a k-nearest neighbour algorithm based on the first 20 PCs and a resolution parameter of 0.6. Clusters were subsequently annotated using canonical gene markers.

### Analysis of synovial cell to cell interactions

To enable a systematic analysis of cell-cell communication molecules, we used CellPhoneDB (Vento-Tormo et al., in press in *Nature*)^23^. CellPhoneDB is a manual curated repository of ligands, receptors and their interactions, integrated with a new statistical framework for inferring cell-cell communication networks from single cell transcriptome data. Briefly, in order to identify the most relevant interactions between cell types, we looked for the cell-type specific interactions between ligands and receptors. Only receptors and ligands expressed in more than 10% of the cells in the specific cluster were considered.

We performed pairwise comparisons between all cell types. We randomly permuted the cluster labels of all cells 1000 times and determined the mean of the average receptor expression level of a cluster and the average ligand expression level of the interacting cluster. For each receptor-ligand pair in each pairwise comparison between two cell types, this generated a null distribution. By calculating the proportion of the means which are “as or more extreme” than the actual mean, we obtained a p-value for the likelihood of cell type-specificity of a given receptor-ligand complex.

### Statistical Analyses

Statistical tests were performed as indicated in the figure legends.

### Data availability

The raw sequencing data generated for the present study has been deposited in the European Genome-phenome Archive using the following study IDs: EGAS00001002701 (Chromium 10X data), EGAS00001002958 (SS2 data). The tables of counts for the 10X (Table S2) and SS2 (Table S3) used in this study have also been made available.

## Acknowledgements

We thank GOSH-ICH Flow Cytometry Facility (Stephanie Channing) for assistance with cell sorting, Dale Godfrey (University of Melbourne), Ildiko Van Rhijn and D. Branch Moody (Harvard Medical School, Boston) for provision of TCR-transduced Jurkat cells and UCL CRUK Cancer Centre Pathology Core Facility for assistance with cytological analysis. We are grateful to Saskia Hemmers, Kylie James and Muzlifah Haniffa for critical review of the manuscript. L.R.W is supported by the NIHR BRC at Great Ormond Street Hospital and Great Ormond Street Children’s Charity. L.R.W, L.M. and M.W. are supported by Arthritis Research UK. Personal fellowships support E.R. (Arthritis Research UK), S.B. (Wellcome; St Baldrick’s Foundation) and C.B. (Wellcome). The views expressed are those of the author(s) and not necessarily those of the NHS, the NIHR or the Department of Health.

## Author Contributions

S.B. and C.B. conceived and designed the experiments; M.D.C.V.H. performed the SMARTtseq2 experiments, designed and performed computational analysis aided by M.D.Y., M.E., R.V.T., L.M.; F.A.V.B. performed the droplet based 10X sequencing, assisted by L.M; E.R., performed flow cytometry and assisted with cell-sorting; C.B. performed flow cytometry experiments and analyzed data; M.W. assisted with sample preparation; H.A.M. performed flow cytometry assays, E.M. performed cytological analysis; A.F. supervised cytological analysis; A.C. assisted with figure preparation, S.T. contributed to discussions; I.V.R. and D.B.M. provided reagents; L.W. provided human samples. S.B. and C.B. wrote the manuscript. L.W., S.B. and C.B. co-directed this study.

## Author Information

The authors declare no conflict of interest. Correspondence and requests for materials should be addressed to.

